# The *Hdc* GC box is critical for *Hdc* gene transcription and histamine-mediated anaphylaxis

**DOI:** 10.1101/2022.06.13.495950

**Authors:** Yapeng Li, Junfeng Gao, Dianzheng Zhao, Xiaoyu Guan, Suzanne C. Morris, Fred D. Finkelman, Hua Huang

## Abstract

**Background**: Histamine is a critical mediator of anaphylaxis, a neurotransmitter, and a regulator of gastric acid secretion. Histidine decarboxylase is a rate-limiting enzyme for histamine synthesis. However, *in vivo* regulation of *Hdc*, the gene that encodes histidine decarboxylase is poorly understood.

**Objective**: We sought to investigate how enhancers regulate *Hdc* gene transcription and histamine synthesis in resting conditions and in a mouse model of anaphylaxis.

**Methods**: H3K27 acetylation histone modification and chromatin accessibility were used to identify candidate enhancers; The enhancer activity of candidate enhancers was measured in a reporter gene assay; and the function enhancers were validated using CRISPR deletion.

**Results**: Deletion of the GC box, which binds to zinc finger transcription factors, in the proximal *Hdc* enhancer, reduced *Hdc* gene transcription and histamine synthesis in the mouse and human mast cell lines. Mast cells, basophils, brain cells, and stomach cells from GC box-deficient mice transcribed the *Hdc* gene much less than similar cells from wild-type mice and *Hdc* GC box-deficient mice failed to develop anaphylaxis.

**Conclusion**: Our results demonstrate that the *HDC* GC box within the proximal enhancer in the mouse and human *HDC* gene is essential for *Hdc* gene transcription, histamine synthesis, and histamine-mediated anaphylaxis *in vitro* and *in vivo*.

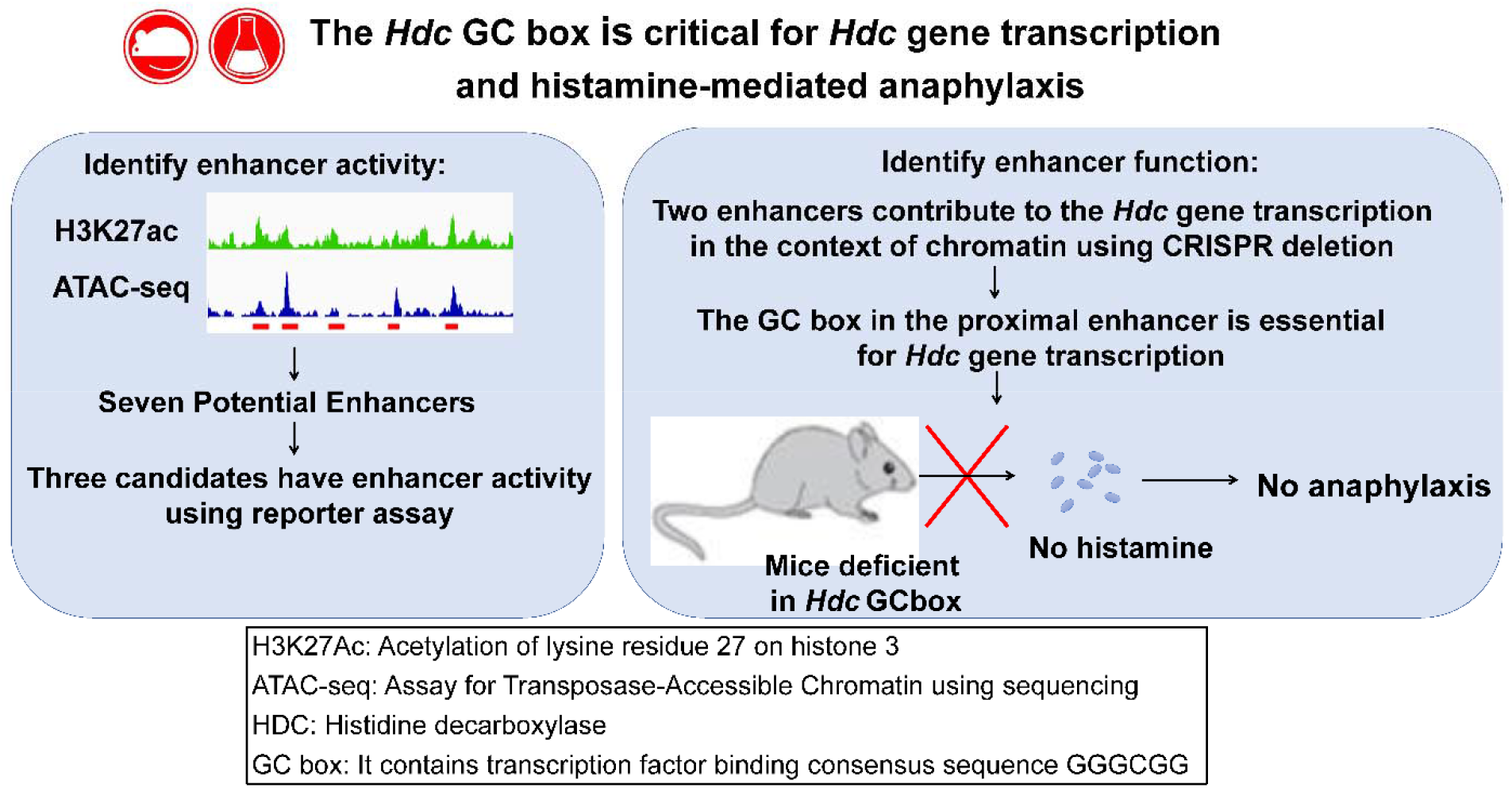

**Key messages**:

- The *HDC* GC box within the proximal enhancer of the mouse and human *HDC* gene is essential for *Hdc* gene transcription and histamine synthesis in mouse and human mast cells.
- The mouse *Hdc* GC box is required for histamine-mediated anaphylaxis *in vivo*.

**Capsule summary**

Our results demonstrate that the *Hdc* GC box within the proximal enhancer of the mouse and human *HDC* gene is essential for *Hdc* gene transcription and histamine synthesis in mast cells. The mouse *Hdc* GC box is required for histamine-mediated anaphylaxis in the mouse model of anaphylaxis.

## Introduction

Anaphylaxis is a life-threatening allergic reaction that rapidly affects the skin, gastrointestinal tract, respiratory system, and cardiovascular system^1^. Anaphylaxis can be caused by allergy to foods, insect venoms, medications, and other agents^1^. Food-induced anaphylaxis has risen dramatically in developed countries during the past several decades and continues to rise^2–4^. Understanding the molecular regulation of anaphylactic shock should promote effective prevention and treatment.

Mast cells (MCs) play a critical role in mediating anaphylaxis by releasing large amounts of histamine and other mediators. MCs express Fc*ε*RI, the high-affinity receptor for IgE, a heterotetramer composed of one IgE-binding α subunit, one membrane-tetraspanning β subunit, and a disulfide-linked γ subunit dimer. MCs can be activated by antigen cross-linking of antigen-specific IgE bound to Fc*ε*RI. Antigen cross-linking of the IgE/Fc*ε*RI complex activates MCs to release histamine that is pre-made and stored in MC granules.

Histamine belongs to a family of biogenic amines that includes neurotransmitters such as serotonin and dopamine and hormones such as adrenaline. Biogenic amines that contain one or more amine groups are formed mainly by decarboxylation of amino acids. Histamine is a monoamine synthesized from the amino acid histidine through a reaction catalyzed by the enzyme histidine decarboxylase (HDC), which removes the carboxyl group from histidine. The *Hdc* gene encodes HDC protein with a molecular mass of 74 kDa, a proenzyme with little or no enzyme activity. However, it becomes active once the proenzyme is cleaved at the site near its c-terminus, presumably by Caspase-9. HDC is the primary enzyme that catalyzes histamine synthesis.

Knowledge of how the human and mouse *Hdc* genes are transcribed is limited^5^. The Mouse *Hdc* gene is located on chromosome 2^6^. It contains 12 exons. *Hdc* mRNA is expressed in basophils, MCs, and enterochromaffin-like cells (ECL) of the stomach and is 86% homologous with the human gene. The human *HDC* gene is located in the 15q21.2 region of chromosome 15 and also contains 12 exons^7^. Most previous work has concentrated on analyzing the *Hdc* promoter region using reporter gene assays in transformed MC cell lines. Deletional analysis of *Hdc* promoter-driven luciferase reporter gene transcription demonstrated that the transcription factor SP1 binds to a GC box (GGGGCGGGG) found in both the human and mouse *Hdc* gene promoters^6, 8^. Several promoter elements have been reported to regulate *Hdc* gene transcription negatively. For example, the transcription factors YY1 and KLF4 negatively regulate the *Hdc* gene by suppressing SP1 in a gastric cancer cell line^9, 10^. Our group demonstrated positive regulation of *Hdc* gene expression by the transcription factor (TF) GATA binding protein 2 (GATA2), a member of the GATA family of TFs and by Microphthalmia-associated TF (MITF)^11^.

Enhancers are segments of DNA located in the noncoding regions of genes. They activate gene transcription by delivering important accessory factors to the promoter^12, 13^. Decoding enhancers has been a longstanding goal in the field of gene transcription^14^. Enhancers and other *cis* regulatory elements work from a distance in eukaryotes. Transcriptional codes hidden in gene distal regions are often required for full transcription. Locating critical enhancers can be difficult because they can be located up to 100 kb from the transcription start sites in noncoding regions, which constitute 99% of the genome. Enhancers contain clusters of TF binding motifs that act as transcriptional codes that are read by TFs. The binding of TFs and co-factors to enhancers creates accessible regions within enhancers that recruit histone-modifying enzymes to add acetyl or methyl groups to histones. Monomethylation of lysine four on histone 3 (H3K4me1) marks genes that are poised to be transcribed, whereas acetylation of lysine 27 on histone 3 (H3K27ac) identifies genes that are actively being transcribed. DNA regions associated with increased histone modification often coincide with chromatin accessibility, which can be measured by ATAC-seq. The combined presence of histone modifications and chromatin accessibility predicts enhancer activity^15–19^ and promotes a looping mechanism that approximates TF- and co-factor-bound distal and proximal enhancers to core promoters via a huge protein complex called mediator^20^; this approximation then facilitates gene transcription. By measuring H3K4me1 and H3K27ac modifications using next generation DNA sequencing, our group has identified a −8.8 kb *Hdc* enhancer that increased minimal *Hdc* promoter activity in a luciferase reporter gene transcription assay^11^.

In this study, by combing H3K27ac modification and chromatin accessibility measurement, we have identified several additional potential enhancers in the 34 kb region of the mouse *Hdc* gene. We have functionally tested the potential enhancers in the context of chromatin in mouse and human mast cell lines and in mice using an improved CRISPR method. Our work demonstrates a critical role for the *HDC* proximal enhancer GC box in human *HDC* gene transcription and histamine synthesis and for murine histamine-mediated anaphylaxis.

## METHODS

### Mice

sgRNA (TTTATGGGGGCGGGGC) was designed and injected into BALB/cByJ zygotes along with ssDNA HDR template. A more detailed description of mice used is included in the Methods section in this article’s Online Repository. All animal experiments were conducted according to protocols approved by the National Jewish Health Institutional Animal Care and Use Committee (Denver, CO).

### BMMC and LAD2 cell culture

BMMCs were cultured from bone marrow cells of BALB/c mice. Human mast cell line LAD2 was provided by Drs. Dean D. Metcalfe and Arnold S. Kirshenbaum (National Institutes of Health, Bethesda, MD). See the Methods section in this article’s Online Repository for further details.

### H3K27ac CUT&Tag-seq and Omni-ATAC-seq

The amplified libraries were purified, size-selected, and the quality and quantity of libraries were assessed with an Agilent Technologies TapeStation. The pair-ended sequencing of DNA libraries was performed with an Illumina NovaSEQ6000 platform. Data can be accessed at the Gene Expression Omnibus database (http://www.ncbi.nlm.nih.gov/geo, GSE223388). See the Methods section in this article’s Online Repository for further details about libraries preparation and NGS data analysis.

### Luciferase reporter constructs and assay

CFTL-15 cells (5 × 10^6^) were electroporated with plasmids using a Bio-Rad Gene Pulser. Luciferase activities were measured by using an InfiniteMlOOO^®^ microplate reader and the Dual-Luciferase reporter assay system. See the Methods section in this article’s Online Repository for further details.

### Deletion of GC boxes using four bicistronic sgRNA guides

The sgRNA sequences targeting *Hdc* enhancers or transcription factor exons were designed using the online CRISPick tool from Broad Institute. Each of four sgRNA sequences targeting the same enhancer was cloned into vector using the BsmBI cloning site. CFTL-15 cells were either transfected with Lenti plasmids or transduced with lentivirus containing the bicistronic sgRNA guides. The sorted cells were subjected to DNA analysis of GC box deletion. The intensities of DNA fragments were measured using Imagej (NIH, MD). See the Methods section in this article’s Online Repository for further details.

### Passive systemic anaphylaxis (PSA)

Mice were sensitized with 10 μg of IgE anti-TNP antibody and challenged with 10 μg of TNP-BSA. Rectal temperatures of the challenged mice were measured before challenge (0 min) and at 5, 10, 15, 20, 25, 30, 40, 60, 80, 100 and 120 minutes after challenge as described previously^11^.

### Histamine and MMCP1 measurements

Plasma was collected 5 minutes after challenge for histamine measurement, using a histamine EIA kit (Beckman Coulter, IM2015). Sera were collected one hour after challenge for determination of serum levels of mucosal mast cell protease (MMCP)1, using an MMCP1 EIA kit (Thermo Fisher Scientific).

## Statistical analysis

The nonparametric Mann-Whitney U test or two-tailed student’s *t* test was used to determine significant differences between two samples. The body temperature changes at various time points before and after PSA challenges were analyzed using a two-way repeated measure ANOVA.

## RESULTS

### The proximal *Hdc* enhancer is essential for *Hdc* gene transcription

Previously, we used the histone modifications H3K4me1 and H3K27ac to identify potential enhancers in the *Hdc* gene and analyzed the distal −8.8 enhancer in detail. In this study, we used H3K27ac CUT&Tag-seq and Omni-ATAC-seq to perform a higher resolution analysis of potential *Hdc* enhancers in a 34 kb region of the *Hdc* gene. In addition to the previously defined *Hdc* E-8.8, we identified proximal enhancer (PE), E+8.5, E+12.3, E+15.8, E+18.8, and E+20.8 noncoding DNA regions that showed increased H3K27ac modification and increased chromatin accessibility (Fig. 1). To determine which of the DNA regions possess enhancer activity, we constructed a series of luciferase reporter genes in which the PE, E+8.5, E+12.3, E+15.8, E+18.8, or E+20.8 was linked to the minimal *Hdc* promoter (MP) (−42 to +1 relative to the TSS). We included a DNA region that was not associated with H3K27ac modification or chromatin accessibility as a negative non-enhancer (NE) control (Fig. 1). The constructs were transfected into the CFTL-15 mast cell line, which we previously used to show *Hdc* E-8.8 enhancer activity^11^. We found that, in addition to *Hdc* E-8.8, the *Hdc* PE and E+18.8 showed enhancer activities relative to the MP promoter (E-8.8, 40.0-fold; PE, 13.4-fold; E+18.8, 12.7-fold) (Fig. 2A). As expected, the NE region did not show enhancer activity (Fig. 2A). These results provide evidence that the *Hdc* PE, E-8.8, and E+18.8 DNA regions contain sites that can bind to cognate transcription factors in CFTL-15 mast cells to promote the transcription of the luciferase reporter gene.

**Figure 1.**
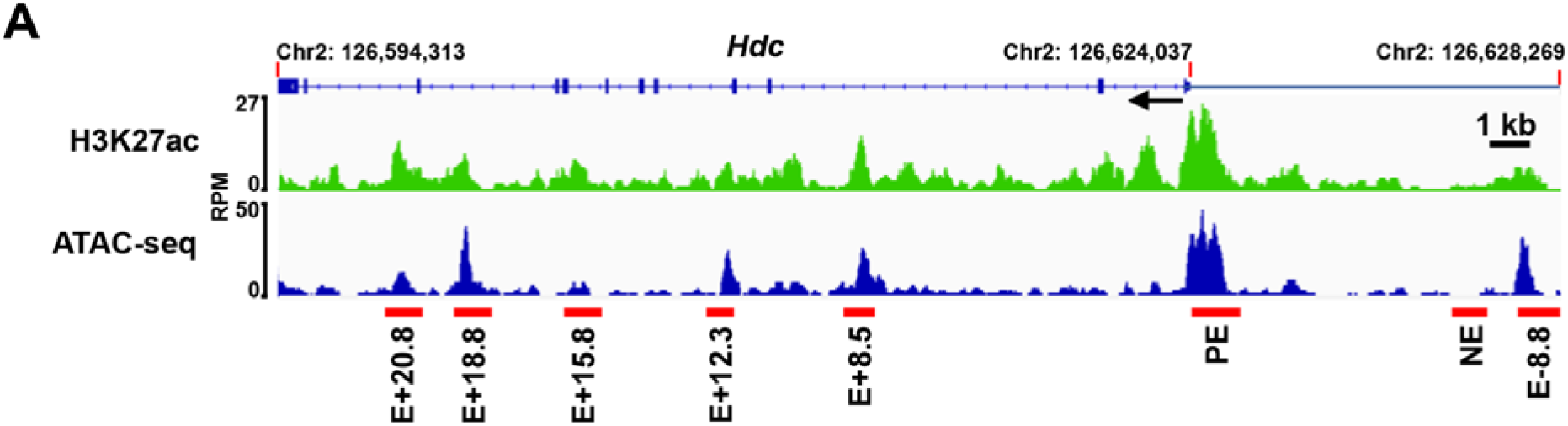
Identification of the putative *Hdc* enhancers. Representative Integrated Genome Viewer (IGV) tracks from H3K27ac CUT&Tag-seq and Omni-ATAC-seq. BMMCs were used for H3K27ac CUT&Tag-seq and Omni-ATAC-seq. Red bars indicate putative *Hdc* enhancers that show increased H3K27ac modifications and chromatin accessibility. RPM: reads per million mapped reads; E: enhancer; PE: proximal enhancer. The numbers following E indicate the distance (kb) of enhancer to the TSS of the *Hdc* gene; + means the enhancers are located downstream of the TSS, and – means the enhancers are located upstream of the TSS. One IGV track was from one biological sample, representing two (Omni-ATAC-seq) or three (CUT&Tag-seq) biological samples with similar patterns.

**Figure 2.**
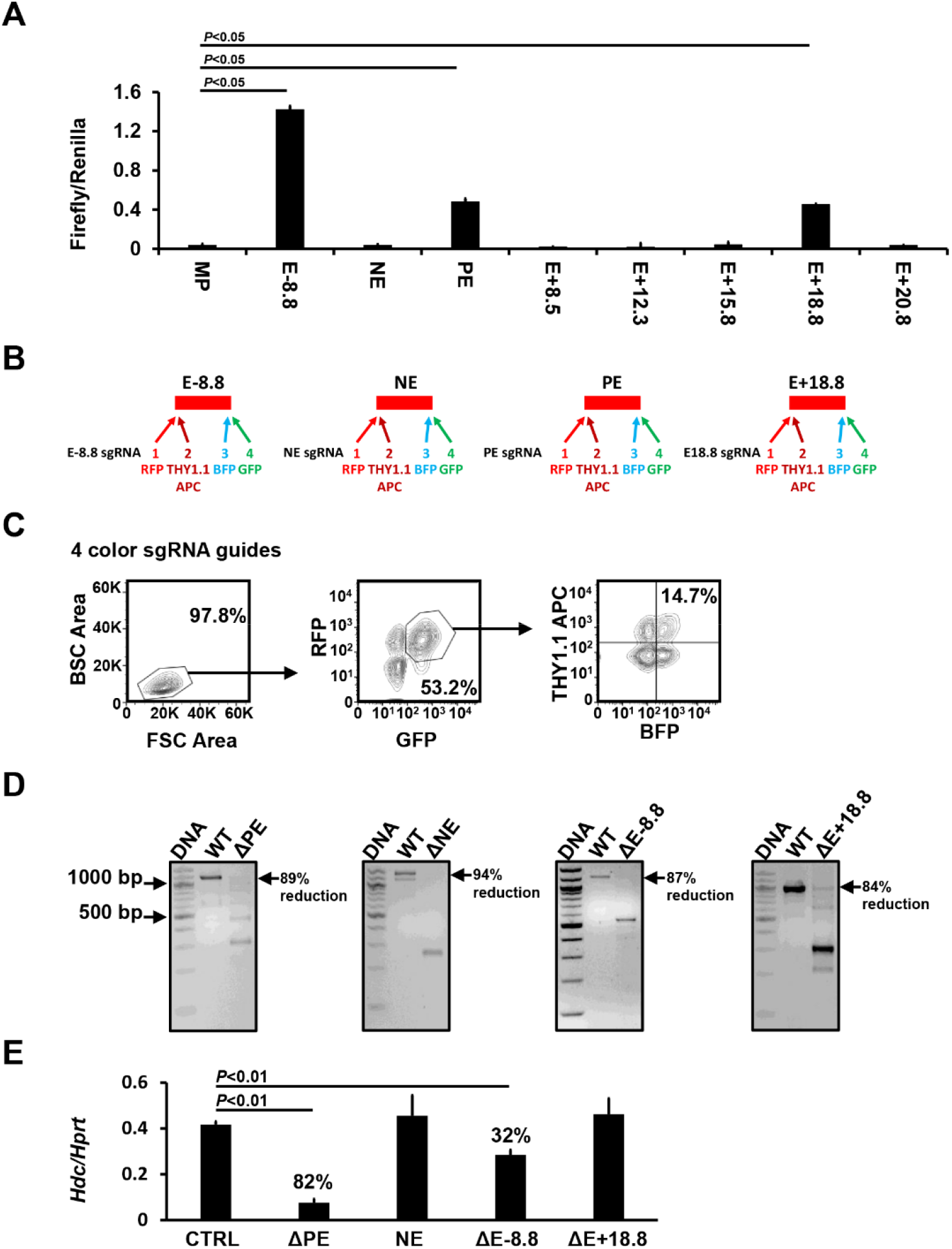
*Hdc* E-8.8, PE, and E+18.8 possess enhancer activity. **A**. Luciferase reporter analysis of the *Hdc* enhancer activities in CFTL-15 mast cells. Error bars represent mean ± SEM, n=4 transfectants from 2 independent experiments. **B**. sgRNA guide constructs that co-expressed bicistronic BFP, GFP, RFP, and Thy1.1 genes. **C**. FACS sorting gates. CFTL15 cells were transduced lentivirus containing four bicistronic sgRNA guides. BFP^+^, GFP^+^, RFP^+^, and Thy1.1^+^ transduced cells were FACS-sorted using sorting gates indicated. **D**. DNA deletion efficiency analysis. WT and deleted DNA fragments in FACS-sorted BFP^+^, GFP^+^, RFP^+^, and Thy1.1^+^ cells were analyzed with PCR. **E**. *Hdc* mRNA expression in the FACS-sorted cells was measured by qPCR (mean ± SEM, n=6 transfectants from 3 independent experiments). The percentages indicate the percent reduction in *Hdc* mRNA expression relative to CTRLs. *P* values were calculated using the Mann-Whitney U test.

However, the enhancer luciferase reporter assay does not assess enhancer activity in the context of chromatin. To determine whether the *Hdc* PE, E-8.8, and/or E+18.8 enhancers play a role in *Hdc* gene transcription in the context of chromatin, we used CRISPR/Cas9 technology to delete these enhancers. We used the NE as a negative control. The CRISPR deletion method using two sgRNA guides often resulted in deletions occurring in one chromosome, creating heterozygous deletion that does not have a phenotype. To overcome this technical challenge, we targeted one enhancer with four sgRNA guides, each expressing a bicistronic gene encoding for fluorescence protein BFP, GFP, RFP, or for Thy1.1 (Fig. 2B). CFTL-15 cells were transduced with lentiviruses containing the four bicistronic sgRNA guides. Cells expressing BFP, GFP, RFP, and Thy1.1 (7.6% of transfected cells were positive for four colors) were FACS-sorted, using gates shown in Fig. 2C. This method achieved 84-94% homozygous deletion of *Hdc* enhancers in bulk (Fig. 2D). Deletion of the PE enhancer resulted in the greatest reduction (82%) in *Hdc* mRNA expression (Fig. 2E), while deletion of the E-8.8 enhancer resulted in a 32% reduction in *Hdc* mRNA expression (Fig. 2E). In contrast, deletion of the NE or *Hdc* E18.8 enhancer did not affect *Hdc* mRNA expression in the context of chromatin (Fig. 2E). These data demonstrate that *Hdc* PE is critical in *Hdc* gene transcription, while the *Hdc* E-8.8 contributes to *Hdc* gene transcription in the context of chromatin.

### The GC box in the proximal *Hdc* enhancer is essential for *Hdc* gene transcription in the CFTL-15 mast cell line

To analyze transcription factor binding sites in the *Hdc* PE enhancer, we performed a deletional analysis of the −1401 bp long *Hdc* PE enhancer using a luciferase reporter gene assay. We found that most PE enhancer activity was driven by sequences located within the −48 bp fragment (Fig. 3A). Transcription factor binding site analysis revealed that a GC box is located within this −48 bp fragment. Mutation in the GC box (GGGCGG, −43 to −48) resulted in a complete loss of enhancer activity in the luciferase reporter gene assay (Fig. 3B). Furthermore, CRISPR-mediated deletion of the GC box in CFTL-15 mast cells reduced *Hdc* mRNA expression by 78% (Fig. 3C). These data demonstrate that the GC box in the proximal *Hdc* PE enhancer is important for *Hdc* gene transcription in a mast cell line.

**Figure 3.**
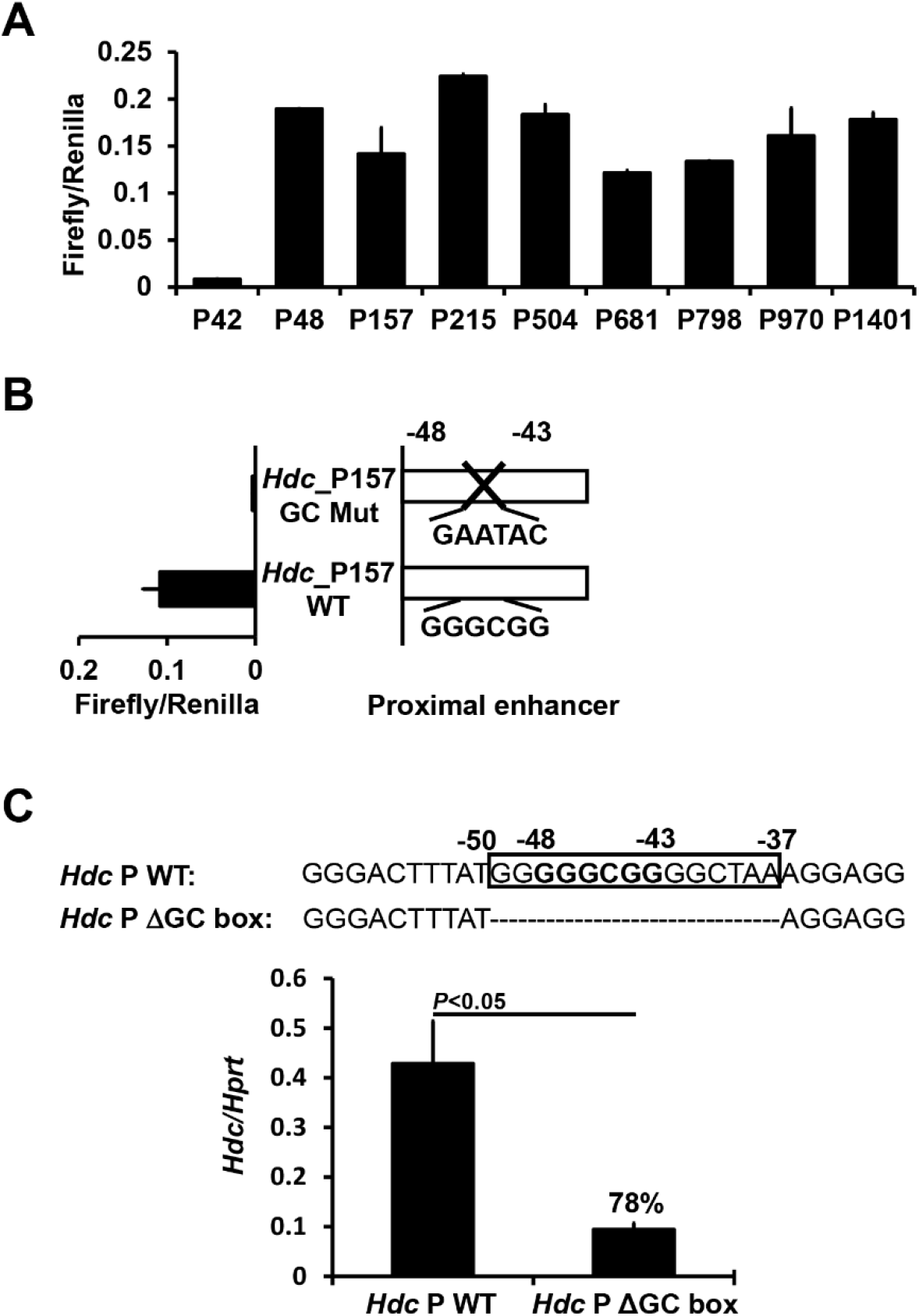
The GC box in the *Hdc* PE is required for *Hdc* gene transcription. **A**. Deletional analysis of the *Hdc* PE in CFTL-15 mast cells. Luciferase reporter genes were transfected into CFTL-15 mast cells. Error bars represent mean ± SEM, n=4 transfectants from 2 independent experiments. **B**. Mutational analysis of the GC box in the −157 *Hdc* PE using a luciferase reporter gene. A luciferase reporter gene containing the GC box mutation was transfected into CFTL-15 mast cells. Mean ± SEMs were calculated from 3 transfectants from 3 independent experiments. **C**. qPCR analysis of *Hdc* mRNA expression in GC box deleted CFTL15 mast cell clones (mean ± SEM, n=3 colonies). CFTL15 mast cells were transfected with the four bicistronic RNA guides. Clones derived from single transfected cells were obtained. The percentages indicate the extent of reduction in *Hdc* mRNA expression. *P* values were calculated using the Mann-Whitney U test.

### Mice deficient in the *Hdc* GC box have greatly reduced synthesis of histamine and IgE-mediated anaphylaxis after antigenic challenge

To validate the findings obtained using the IL-3-dependent non-transformed mast cell line CFTL-15, we generated mice deficient in the *Hdc* GC box using a single sgRNA guide as shown in Fig. 4A to make double-stranded breaks and a homology-directed repair template that lacks the GC box to repair the breaks. A founder strain of mice with a six-nucleotide deletion of the *Hdc* GC box was verified by sequencing (Fig. 4A). Mice deficient in the *Hdc* GC box (*Hdc*^GCbox−/−^) tended to have slightly reduced numbers of peritoneal mast cells (Fig. E1A and B). Surprisingly, Fc*ε*RIα expression, but not cKit expression, on the peritoneal mast cells in the *Hdc*^GCbox−/−^ mice tended to be elevated compared to control *Hdc*^GCbox+/+^ mice (Fig. E1C). To determine whether the tendency towards increased peritoneal mast cell Fc*ε*RIα expression was a direct or indirect consequence of the *Hdc* GC box deletion, we compared Fc*ε*RIα and cKit expression by BMMCs derived from *Hdc*^GCbox+/+^ and *Hdc*^GCbox−/−^ mice *in vitro*, where the indirect extrinsic interactions that occur *in vivo* are reduced. We found a noticeable difference in Fc*ε*RIα expression but not cKit expression on *Hdc*^GCbox−/−^ BMMCs relative to *Hdc*^GCbox+/+^ BMMCs. A small population of BMMCs expressing relatively low levels of Fc*ε*RIα was observed in *Hdc*^GCbox+/+^ but not *Hdc*^GCbox−/−^ BMMCs (Fig. E2). The difference may be noticeable but it is not statistically significant. In contrast, *Hdc* mRNA expression was considerably and significantly reduced in both BMMCs (90% reduction) and basophils (77% reduction) (Fig. 4B). To determine whether the *Hdc* GC box only regulates the transcription of the *Hdc* gene, we examined mRNA expression of a neighboring gene *Gabpb1*, which is located just 10 kb upstream of the *Hdc* gene. Deletion of the *Hdc* GC box did not affect *Gabpb1* mRNA expression, supporting that the GC box in the PE of the *Hdc* gene acts explicitly on the transcription of the *Hdc* gene (Fig. 4B).

**Figure 4.**
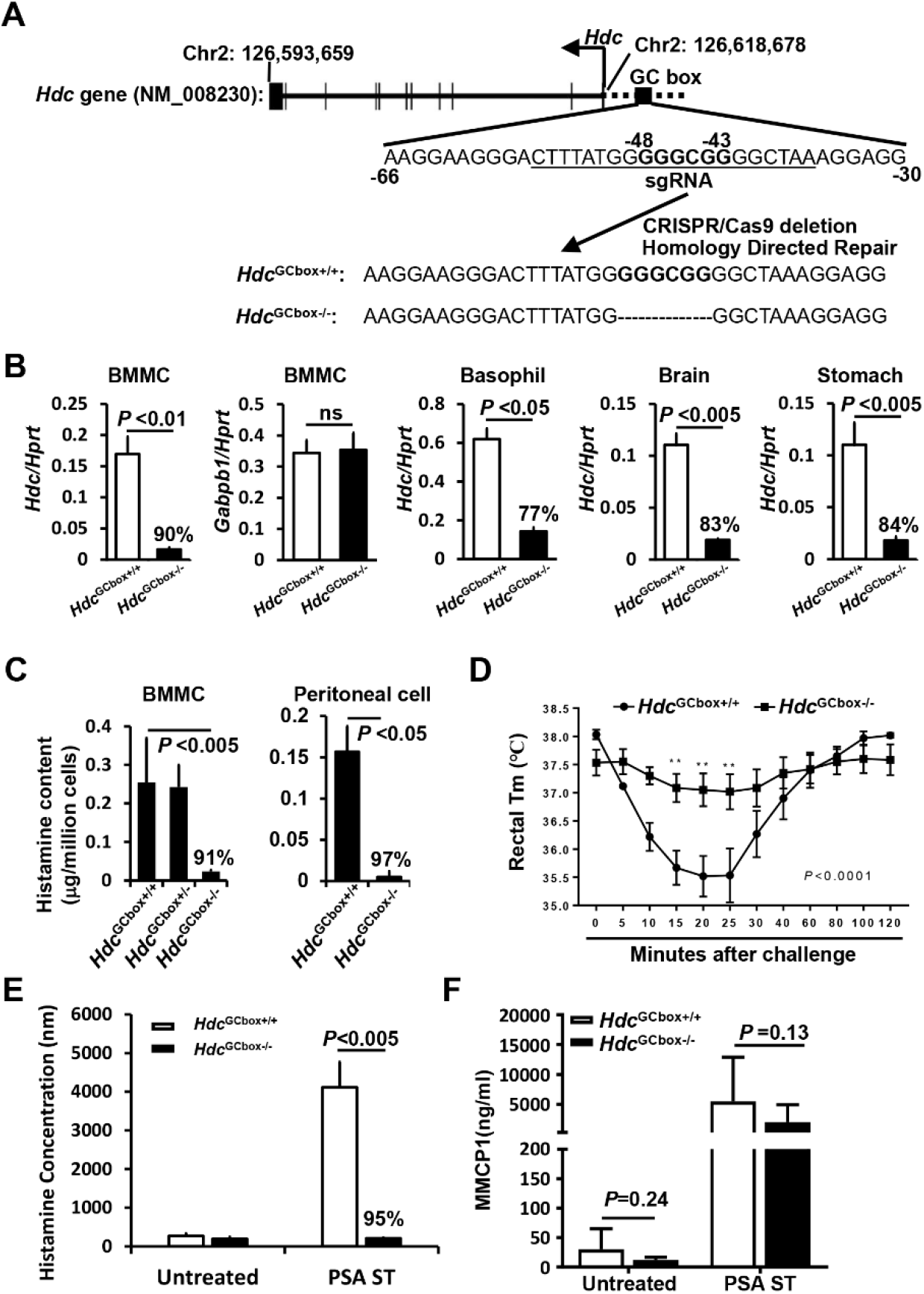
The GC box in the *Hdc* PE is critical for histamine synthesis and IgE/mast cell-mediated anaphylaxis. **A**. mice deficient in GC box (GGGCGG) were generated by a single sgRNA (underlined) targeting the GC box located from-48 to −43 in the proximal promoter of the *Hdc* gene. **B**. qPCR analysis of *Hdc* and *Gabpb1* mRNA expression in cells isolated from *Hdc*^GCbox+/+^ and *Hdc*^GC−/−^ mice (n=4, pooled from two experiments). **C**. histamine contents in BMMCs derived from *Hdc*^GCbox+/+^ and *Hdc*^GC−/−^ mice (n=6, two experiments). **D**. PSA of *Hdc*^GCbox+/+^ and *Hdc*^GC−/−^ mice (n=6, two experiments). **E**. histamine release in the plasma of *Hdc*^GCbox+/+^ and *Hdc*^GC−/−^ mice that were treated with IgE anti-TNP antibody followed by challenge with BSA-TNP. Blood was drawn five minutes after challenge (n=6, two experiments). **F**. MMCP1 concentrations in the sera of *Hdc*^GCbox+/+^ and *Hdc*^GC−/−^ mice that were treated with IgE anti-TNP antibody and BSA-TNP for one hour (n=6, two experiments). *P* values were calculated using the Mann-Whitney U test.

In addition to its activity as an allergic mediator, histamine is a neurotransmitter and a regulator of gastric acid secretion^5^. Histaminergic neurons in the basal ganglia of the brain express *Hdc* mRNA and synthesize histamine. In the stomach, enterochromaffin-like cells express *Hdc* mRNA and synthesize histamine, which stimulates parietal cells in the stomach to secrete stomach acid^21, 22^. We found that the levels of *Hdc* mRNA were significantly decreased in brain cells (83% reduction) and stomach cells (84% reduction) in the *Hdc*^GCbox−/−^ mice (Fig. 4B). Consistent with the reduction of *Hdc* mRNA in the *Hdc*^GCbox−/−^ mice, histamine contents in BMMCs and peritoneal mast cells derived from the *Hdc*^GCbox−/−^ mice were dramatically decreased (91% and 97% reduction, respectively, Fig. 4C).

Histamine plays a critical role in IgE/mast cell-mediated anaphylaxis. We found that the *Hdc*^GCbox−/−^ mice developed little or no passive systemic anaphylaxis (Fig. 4D) and released little histamine (Fig. 4E) after the injection of IgE anti-TNP antibody and challenge with TNP-BSA (95% reduction in histamine release relative to *Hdc*^GCbox+/+^ mice). In contrast, mast cell protease MMCP-1 release to the blood was reduced moderately, if at all, in *Hdc*^GCbox−/−^ mice compared to *Hdc*^GCbox+/+^ mice (Fig. 4F), indicating that the *Hdc* GC box deletion primarily affects histamine synthesis. These data demonstrate that the GC box in the proximal *Hdc* enhancer is critical for histamine synthesis and histamine-mediated anaphylaxis *in vivo*.

### The GC box plays a minor role in regulating chromatin modification and chromatin accessibility at the distal enhancers of the *Hdc* gene

Because distal enhancers can interact with proximal enhancers and core promoters through a looping mechanism, it is possible that the lack of an *Hdc* GC box in the proximal enhancer could affect enhancer structures of distal enhancers^16^. For this reason, we investigated whether the *Hdc* GC box regulates chromatin structure at the distal enhancers. We performed Omni-ATAC-seq and H3K27ac ChIP-seq in *Hdc*^GCbox+/+^ and *Hdc*^GCbox−/−^ BMMCs. Chromatin accessibility at the PE enhancer near the GC box was notably reduced in *Hdc*^GCbox−/−^ BMMCs. Reads at two out of three peaks (peak one and two) near the deleted GC Box were reduced by 65% and 43%, respectively (Fig. 5A). However, H3K27ac histone modification was not reduced by the absence of the *Hdc* GC box. Instead, the H3K27ac peak at the −8.8 position was elevated (Fig. 5B), indicating that *the Hdc* GC box plays a minor role in chromatin accessibility at the PE enhancer and does not contribute to chromatin accessibility at the distal enhancers. These data demonstrate that the *Hdc* GC box plays a minor role in regulating chromatin modification and chromatin accessibility at distal enhancers of the *Hdc* gene.

**Figure 5.**
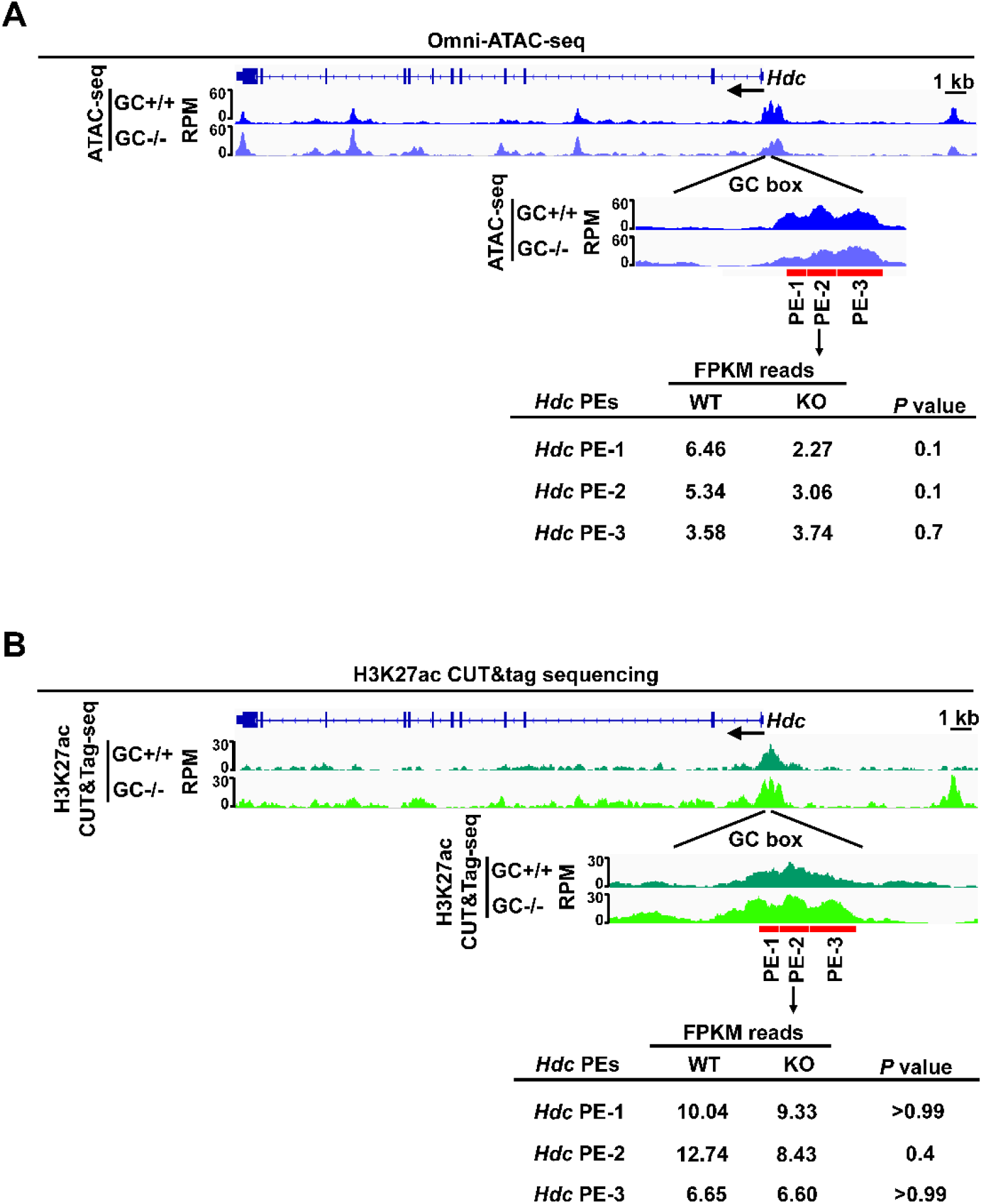
GC box only promotes chromatin accessibility, but not H3K27ac modification, at the *Hdc* PE. **A**. Representative Integrated Genome Viewer (IGV) tracks from Omni-ATAC-seq (n=3). **B**. Representative Integrated Genome Viewer (IGV) tracks from H3K27ac CUT&Tag-seq representing three biological samples (n=3). Statistical significance was analyzed with a two-tailed student’s *t* test.

### The GC box is required for *HDC* gene transcription and histamine synthesis in human mast cells

To determine whether human *HDC* is also regulated by the GC box in the context of chromatin, we designed four bicistronic RNA guides targeting the human *HDC* GC box in the proximal enhancers (−153 to −495, Fig. 6 A). Human mast cell line LAD2 cells were transduced with lentivirus containing the four bicistronic sgRNA guides. LAD2 cells expressing BFP, GFP, RFP, and Thy1.1 (17.6% of transfected cells were positive for 4 colors) were FACS-sorted, using gates shown in Fig. 6B. We achieved 91% homozygous deletion of the GC box (Fig. 6C). Deletion of the *HDC* GC box resulted in 77% reduction in *HDC* mRNA expression (Fig. 6D). Consistent with this, histamine content in the GC box-deleted LAD2 cells was reduced by 86% (Fig. 6E). These results demonstrate that the human *HDC* GC box is as critical for *HDC* gene transcription in human MCs as in mouse MCs.

**Figure 6.**
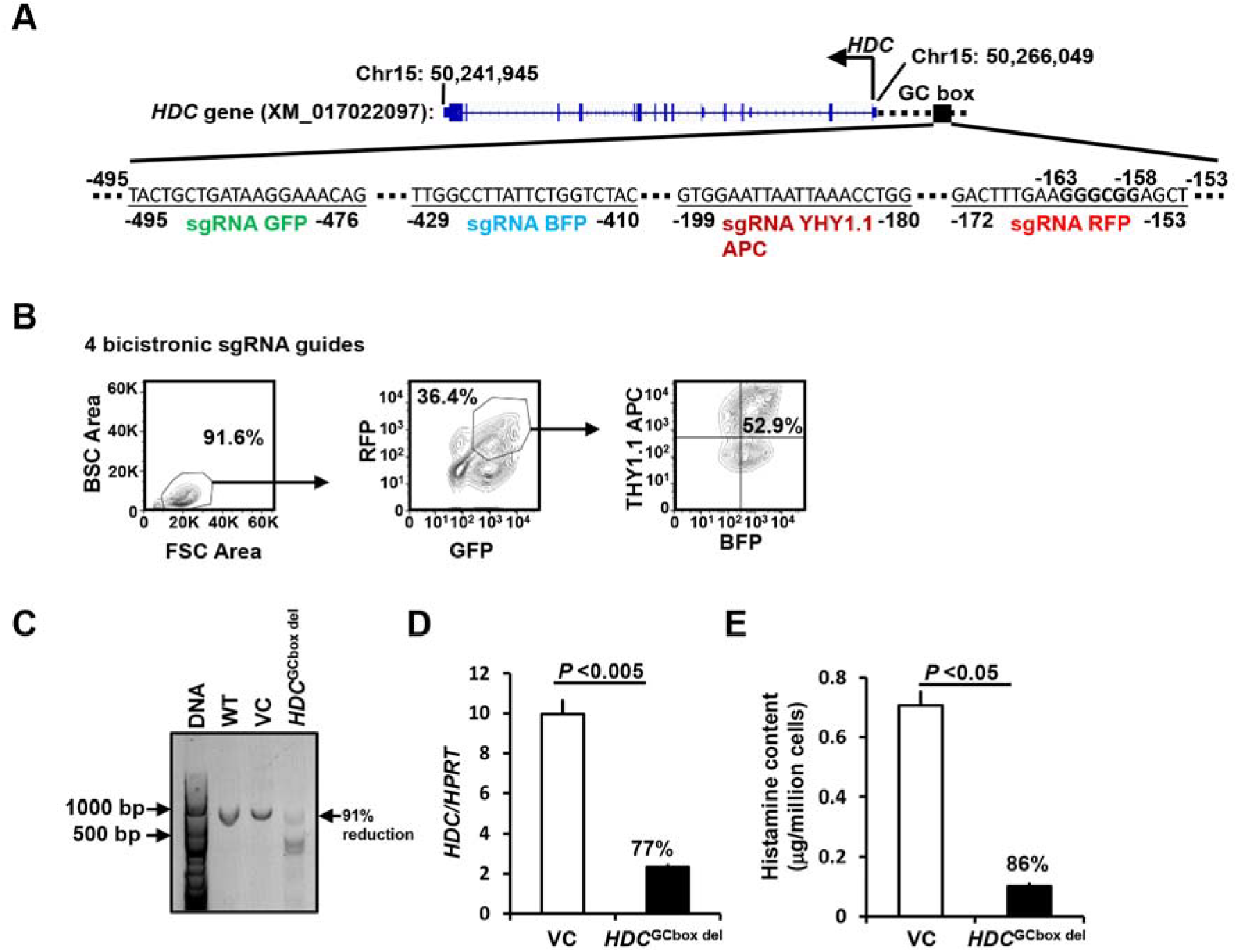
The *HDC* GC box in the human *HDC* gene is critically important for *HDC* gene transcription and histamine synthesis. **A**. Bicistronic BFP, GFP, RFP, and Thy1.1 sgRNA guides targeting the human *HDC* GC box located in −153 to −495. **B**. FACS sorting gates. LAD2 cells were transduced with lentivirus containing four bicistronic sgRNA guides. BFP^+^, GFP^+^, RFP^+^, and Thy1.1 APC^+^ transfectant cells were FACS-sorted using the sorting gates indicated (Fig. 2A). **C**. DNA deletion efficiency analysis. GC box fragments from untransduced WT, or FACS-sorted BFP^+^, GFP^+^, RFP^+^, and Thy1.1APC^+^ cells transduced with lenti vector without sgRNA guide vector controls (VCs) and with four sgRNA guides targeting the human *HDC* GC box were analyzed with PCR. Percentage of reduction was calculated by reduction in density of the WT band. **D**. *HDC* mRNA expression in the FACS-sorted BFP^+^, GFP^+^, RFP^+^, and Thy1.1APC^+^ cells was measured by qPCR (mean ± SEM, n=4 from 2 independent experiments). **E**. Histamine content in the FACS-sorted BFP^+^, GFP^+^, RFP^+^, and Thy1.1APC^+^ cells was measured by ELISA (mean ± SEM, n=4 from 2 independent experiments). The percentages indicate the percent reduction in *HDC* mRNA expression and histamine content relative to VCs. *P* values were calculated using the Mann-Whitney U test.

## DISCUSSION

The work described in this paper contributes to the study and understanding of eukaryote gene transcription in general and *Hdc* gene transcription in particular in several ways. It: (1) moves beyond *in vitro* studies with luciferase reporter genes to *in vivo* studies that incorporate the effects of chromatin; (2) reveals and validates an improved method for CRISPR-mediated homozygous deletion of a gene segment; (3) demonstrates a critical role for the proximal enhancer GC box in both mouse and human *HDC* gene expression despite its lack of influence on the chromatin structure of *Hdc* distal enhancers; and (4) provides a novel and useful mouse model in which nearly absent *Hdc* transcription blocks histamine synthesis and IgE-mediated systemic anaphylaxis with little or no reduction in mast cell number, maturation, or degranulation.

Genes are composed of coding and noncoding DNA regions, spanning tens of kbs in length. Enhancers are noncoding DNA regions that contain cis regulatory regions that are bound by transcription factors and co-factors, such as histone modifying enzymes that can add and remove acetylated or methylated lysines in the histones. Enhancers are often surrounded by chromatin that has histones with acetylated lysines that make the enhancers more accessible to transcription factors and co-factors. The traditional method of analyzing enhancer activity transfects cell lines with reporter genes; although this is a powerful approach for predicting enhancer activity, it has the limitation of ignoring the role of chromatin in enhancer function.

The remarkable progress that has been made in CRISPR gene editing technology now enables analysis of enhancer activity in the context of chromatin by deleting enhancers *in situ*. However, the typical CRISPR deletion method, which uses two sgRNA guides, often deletes the targeted gene in only one chromosome, producing heterozygotes that lack a phenotype. To overcome this technical challenge, we have used four sgRNA guides, each expressing a bicistronic gene that encodes Thy1.1 or a fluorescence protein (BFP, GFP, or RFP), to target an enhancer. By coupling this with fluorescence activated cell sorting to select for cells that express all four marker genes, we achieved 84% to 94% enhancer deletion at both chromosomes without having to perform single-cell cloning. This technical improvement has allowed us to analyze mouse and human *HDC* gene transcription in the context of chromatin. Prior to the development of this technique, we used a luciferase reporter gene assay with the murine CFTL-15 MC cell line to demonstrate that the distal *Hdc* E-8.8 enhancer contributes to *Hdc* gene transcription. Now, using a novel sgRNA technique, we have demonstrated that this enhancer contributes to mouse *Hdc* gene transcription in the context of chromatin. Furthermore, our work has revealed a critical role of the GC box in both mouse and human *HDC* gene in the context of chromatin.

The GC box has a consensus sequence of GGGCGG and is often located near the promoter of eukaryotic genes. Members of the 3-zinc finger TF family, which includes SP1, Krox/Egr, Wilms’ tumor, MIGI, and CREA, can bind to the GC box (GGGCGG)^23^, where they function similarly to TFs that bind to more distal enhancers. There is only a single GC box in the mouse and human *HDC* promoters, although most promoters that express the GC box express it multiple times^24^. Remarkably, deletion of the single GC box in the mouse or human *HDC* proximal enhancer in the context of chromatin profoundly reduces *Hdc* gene transcription. The importance of the mouse *Hdc* GC box in allergy is demonstrated further by our study of its role in a mouse model of anaphylaxis. This validation of the GC box function in mice adds rigor and greater biologic relevance to previously published conclusions, which depended on studies with a single cell line that had been generated from a single mouse strain and used a mast cell line that had been cultured for a long time. Our *in vivo* validation allows broader generalization of our finding that the *Hdc* GC box is required for *Hdc* gene transcription, histamine synthesis, and histamine-mediated anaphylaxis. Currently, it is not clear which members of 3-zinc finger TF family bind to the mouse and human *HDC* GC box under the resting and antigen-stimulated conditions.

The functioning of distal enhancers, proximal enhancers, and core promoters through a looping mechanism^20^ makes it possible that modifying or eliminating the GC box in proximal enhancer could result in profound changes in enhancer structures at the distal enhancers.

However, despite its importance in *Hdc* gene transcription, absence of the GC box in the proximal *Hdc* enhancer slightly alters and does not reduce the chromatin structures of distal enhancers of the mouse *Hdc* gene. This is consistent with the view that other transcription factors, such as GATA2, control *Hdc* gene chromatin accessibility^11^. The GC box thus might function as an enhancer to promote the assembly of the transcriptional complex.

In contrast to the existing *Hdc* gene knockout mice^25^, the *Hdc*^GCbox−/−^ mice that we have generated show only minor effects on mast cell maturation. The number of peritoneal cavity mast cells, which are considered mature mast cells, appears to be slightly, but not significantly reduced. Surprisingly, cKIT and Fc*ε*RIα on *Hdc*^GCbox−/−^ peritoneal cavity mast cells were expressed at normal or higher levels than in *Hdc*^GCbox+/+^ mast cells. Consistent with the notion that the *Hdc* GC box primarily regulates *Hdc* gene transcription, serum mast cell protease MMCP1 levels in naive *Hdc*^GCbox−/−^ mice and *Hdc*^GCbox−/−^ mice that had undergone passive systemic anaphylaxis were reduced moderately, if at all. Any reduction might have been caused by a proportionate reduction in the number of mature mast cells in *Hdc*^GCbox−/−^ mice. Because *Hdc*^GCbox−/−^ mice have relatively normal mast cell differentiation, these mice are more suitable than global *Hdc*-deficient mice for studying the importance of the *Hdc* gene in mast cell/IgE-mediated anaphylaxis. Additionally, because *Hdc*^GCbox−/−^ mice also have reduced *Hdc* gene transcription in stomach and brain tissues, these mice can be used to study how the *Hdc* gene is regulated in neurons and in gastric enterochromaffin-like cells.

## Supporting information

Online Repository

## Abbreviations used

BMMCs: Bone marrow-derived mast cells
ATAC-seq: Assay for Transposase-Accessible Chromatin using sequencing
CUT&Tag-seq: The Cleavage Under Targets and Tagmentation
H3K27Ac: Acetylation of lysine residue 27 on histone 3
FACS: Fluorescence activated cell sorting
GC box: It contains transcription factor binding consensus sequence GGGCGG
HDC: Histidine decarboxylase
PCMCs: Peritoneal cavity mast cells
MMCP1: Mucosal mast cell protease-1
PSA: Passive systemic anaphylaxis
TNP: 2, 4, 6-Trinitrophenyl
TSS: Transcription start site

## Acknowledgements

We thank lab members for thoughtful discussions. We thank Ms. Marlene Gallegos-Sanchez, Kirby Motsinger for technical assistance. We are grateful to Dr. Melissa Brown of Northwestern University for providing us with the CFTL-15 mast cell line; to Drs. Dean Metcalfe and Arnold S. Kirshenbaum of National Institute of Allergy and Infectious Diseases for providing us with the human mast cell line LAD2. We thank Dr. Jennifer Matsuda and her staff in the Regional Mouse Genetics Core Facility at National Jewish Health for generating mice deficient in GC box. We thank the staff at the Animal Care Facility for animal care and other technical assistance. We thank Josh Loomis and Shirley Sobus of the Cytometry Core cell sorting and FACS analysis. We thank staff at genomic core facility at University of Colorado and the Gene and Environment Center at National Jewish Health for NGS sequencing.

## Author contributions

Conceptualization, H.H., Y.L. and F.D.F.; methodology, H.H., Y.L., J.G., D.Z., S.C.M and F.D.F.; investigation, Y.L., J.G., D.Z., X.G. and S.C.M; visualization, Y.L., J.G., D.Z. and X.G.; funding acquisition, H.H. and F.D.F.; supervision, H.H. and F.D.F.; writing-original draft, H.H. and Y.L.; writing-review & editing, H.H., Y.L., F.D.F., X.G., S.C.M, J.G. and D.Z.

