## Supplementary material for "The *Hdc* GC box is critical for *Hdc* gene transcription and histamine-mediated anaphylaxis": Online Repository

**METHODS**

**Mice**

sgRNA (TTTATGGGGGCGGGGC) was designed using the online CRISPick {Doench, 2016 #5840}{Sanson, 2018 #5841}tool from Broad Institute (<https://portals.broadinstitute.org/>) and was injected into BALB/cByJ zygotes along with ssDNA HDR template (CAAATCAGAAAGGACGGGGACAAGAGAGGCTGAAATTAAGCCTGAAGGAAGGGACTTTATGGGGCTAAAGGAGGGGAATAGGGAACTCCTTAAATAAGAGATGCACTGGCTGCCAGGGAGTGCAC) as described^E1^. BALB/cByJ mice (Stock No. 001026) were purchased from the Jackson Laboratory (Bar Harbor, ME). The six-nucleotide deletion (GGGCGG) in *Hdc*^GCbox+/-^mice was verified by sequencing. Two out of eleven founders that have an exact GGGCGG six-nucleotide deletion were selected for further breeding. Mice deficient in the GC box were backcrossed to BALB/c mice for one generation and designated as Balb/c-Hdc^em1huhu^/J mice. These mice are available from the Jackson Laboratory (Stock No. 037608). Genotyping was carried out using the following primers: forward, TCAAATCAGAAAGGACGGGGA; reverse, ATTTAAGGAGTTCCCTATTCCCCTC.

**BMMC and LAD2 cell culture**

Bone marrow-derived mast cells (BMMCs) were cultured from bone marrow cells of BALB/c mice in Iscove’s DMEM (10016CV, Corning^TM^ cellgro^TM^, Manassas, VA) plus 10% FBS,100 units/mL penicillin, 100 μg/mL streptomycin, and 2 mM β-mercaptoethanol in the presence of 20 ng/mL IL-3 for four weeks. Over 99% of BMMCs were mast cells had a mast cell phenotype by FACS analysis (FcεRIα^+^ c-Kit^+^). Human mast cell line LAD2 was provided by Drs. Dean D. Metcalfe and Arnold S. Kirshenbaum (National Institutes of Health, Bethesda, MD) and cultured in AIM V™ Medium (Cat#31035025, Thermo Fisher Scientific, Waltham, MA) plus 2 mM beta-mercaptoethanol, 2 mM L-Glutamine (Cat#25005CI, Thermo Fisher Scientific, Waltham, MA), 100 units/ml penicillin, and 100 µg/ml streptomycin in the presence of 100 ng/ml recombinant human SCF (Cat#300-07, Peprotech, Cranbury, NJ).

**H3K27ac CUT&Tag-seq**

The Cleavage Under Targets and Tagmentation (CUT&Tag) assay for H3K27ac modification was performed according to the published protocol^E2^. Omni-ATAC-seq was performed as previously described^E3, 4^. BMMCs (1 × 10^5^) were treated with concanavalin A-coated beads (Bangs Laboratories, BP531) and incubated with rabbit anti-H3K27ac antibody (Abcam, ab4729) at 4 ℃ overnight. The cells were then incubated with guinea pig anti-rabbit antibody (Antibodies online, ABIN101961) at room temperature for 30 minutes. The cells were washed and incubated with pA-Tn5 adapter complex (Diagenode, C01070001) at room temperature for 1 hour on a rotator. The cell pellet was resuspended in 300 μl tagmentation buffer and incubated at 37 ℃ for 1 hour. The reaction was stopped by adding 10 μL 0.5M EDTA, 3 μL 10% SDS, and 2.5 μL 20 mg/mL Proteinase K, and cells were digested at 37 ℃ overnight. Then, DNA was extracted using phenol chloroform and diluted in 25 μl TE buffer with 1/400 RNAse A. The tagmented DNA was enriched by PCR amplification using Universal i5 primer and uniquely barcoded i7 primers under the following conditions: 72 °C for 5 min, and 13 cycles: 98 °C for 30 sec, 98 °C for 10 sec, 63 °C for 10 sec, 72°C for 1 min and hold at 8 ℃. The PCR products were cleaned up and size-selected using Ampure XP beads (Beckman Coulter, A63880). The quality and quantity of libraries were analyzed with the Agilent TapeStation 4200 system. Pair-ended sequencing of DNA libraries was performed on an Illumina NovaSEQ6000 platform.

**Omni-ATAC-seq**

BMMCs (5 × 10^4^) either untreated or activated by IgE receptor crosslinking, were spun down and washed once with cold PBS. The cells were resuspended in 50 μl cold ATAC-RSB-lysis buffer (10mM Tris-HCl pH 7.4, 10mM NaCl, 3mM MgCl_2_, 0.1% NP-40, 0.1% Tween-20 and 0.01% digitonin) and incubated for 3 minutes. The lysis buffer was immediately washed out with 1 mL ATAC-RSB buffer (10mM Tris-HCl pH 7.4, 10mM NaCl, 3mM MgCl_2_ and 0.1% Tween-20). The cell pellet was resuspended in 50 μl transposition mix (25 μl 2X TD buffer, 2.5 μl transposase (Illumina, FC-121-1030), 16.5 μl PBS, 0.5 μl 1% digitonin, 0.5 μl 10% Tween-20, 5 μl H_2_O) and incubated at 37 °C for 30 minutes. The reaction was stopped by adding 2.5 μl of 0.5M EDTA, pH 8, and transposed DNA was purified with a Qiagen MiniElute PCR purification kit (Qiagen). Purified DNA was amplified using the following conditions: 72°C for 5 min, 98 °C for 30 s, and 7 cycles: 98 °C for 10 s, 63 °C for 30 s, 72 °C for 1 min. The amplified libraries were purified, size-selected, and the quality and quantity of libraries were assessed with an Agilent Technologies 2100 Bioanalyzer. The pair-ended sequencing of DNA libraries was performed with an Illumina NovaSEQ6000 platform.

**NGS data analysis**

Raw sequencing reads (average 118.2 million reads, 2 biological replicates) were adaptor trimmed by Trimmomatic 0.33 and the quality of sequenced data was analyzed by FastQC. The trimmed reads were aligned to the mm10 reference genome using Bowtie2 with -very-sensitive and -x 2000 parameters. The read alignments were filtered using SAMtools (version 1.7) to remove mitochondrial genome and PCR duplicates. Peaks were identified by MACS2 with the q-value cut-off of 0.05 and the sequencing data was displayed using IGV. The representative ATAC-seq IGV tracks showed in the figures were generated from one biological sample, representing two or three biological samples with similar pattens.

**Luciferase reporter constructs and assay**

CFTL-15 cells (5 × 10^6^) were electroporated with 10 μg of luciferase plasmid and 0.5 μg of *Renilla* vector at 450 V and 500 microfarads using a Bio-Rad Gene Pulser. Twenty hours after electroporation or transfection, cells were collected, and luciferase activities were measured by using an InfiniteMlOOO® microplate reader (Tecan Systems, Inc., San Jose, CA) and the Dual-Luciferase reporter assay system (E1960, Promega). Transcriptional activity was normalized as the ratio of luciferase activity divided by *Renilla* activity.

**Deletion of GC boxes using four bicistronic sgRNA guides**

The sgRNA sequences targeting *Hdc* enhancers or transcription factor exons were designed using the online CRISPick tool from Broad Institute (<https://portals.broadinstitute.org/>). Each of four sgRNA sequences targeting the same enhancer was cloned into LentiCRISPRv2GFP (Addgene, Plasmid # 82416), LentiCRISPRv2-mCherry (Addgene, Plasmid #99154), LentiCRISPRv2BFP, or LentiCRISPRv2THY1.1 vectors. LentiCRISPRv2THY1.1 vector was modified by replacing the *Gfp* gene in LentiCRISPRv2GFP with the gene encoding THY1.1, using the BsmBI cloning site. CFTL-15 cells were either transfected with Lenti plasmids or transduced with lentivirus containing the bicistronic sgRNA guides. For transduction, lentiviruses were prepared using method describe in our publication. CFTL-15 mast cells were transduced using spin infections as described; For transfection, CFTL-15 cells (10 × 10^6^) were electroporated with 10 μg of each CRISPR plasmid using a Bio-Rad Gene Pulser at 450 V and 500 microfarads. Twenty-four hours after electroporation, the cells that expressed BFP, GFP, RFP, and Thy1.1 were FACS-sorted. The sorted cells were subjected to DNA analysis of GC box deletion. The intensities of DNA fragments were measured using Imagej (NIH, MD).

**Figure Legends**

**FIG E1. Mice deficient in the *Hdc* GC box had slightly reduced numbers and expressed higher FcεRIα and cKit expression levels in peritoneal mast cells**

**A**. Peritoneal cells were collected from WT and *Hdc* GC box (*Hdc*^GCbox-/-^) mice and analyzed by FACS, DAPI^-^ CD3ε^-^ B220^-^ FcεRIα ^+^ c-Kit^+^ peritoneal mast cells were shown in the gates. **B**. Percentage and total numbers of peritoneal mast cells in the WT and *Hdc* GC box (*Hdc*^GCbox-/-^) mice (mean ± SEM, n=4 from 2 independent experiments). **C**. MFI of FcεRIα and cKit of the peritoneal mast cells in the WT and *Hdc* GC box (*Hdc*^GCbox-/-^) mice (mean ± SEM, n=4 from 2 independent experiments). *P* values were calculated using the Mann-Whitney U test.

**FIG E2. Mice deficient in the *Hdc* GC box lack a population of BMMCs that have low FcεRI expression.** **A**. BMMCs were cultured from WT and *Hdc* GC box (*Hdc*^GCbox-/-^) mice and analyzed by FACS using gates shown. **B**. MFI of FcεRIα and c-Kit on WT and *Hdc*^GCbox-/-^ BMMCs (mean ± SEM, n=3 from 2 independent experiments). *P* values were calculated using the Mann-Whitney U test.
